# Measuring the invisible – The sequences causal of genome size differences in eyebrights (*Euphrasia*) revealed by k-mers

**DOI:** 10.1101/2021.11.09.467866

**Authors:** Hannes Becher, Jacob Sampson, Alex D. Twyford

## Abstract

Genome size variation within plant (and other) taxa may be due to presence/absence variation in low-copy sequences or copy number variation in genomic repeats of various frequency classes. However, identifying the sequences underpinning genome size variation has been challenging because genome assemblies commonly contain collapsed representations of repetitive sequences and because genome skimming studies miss low-copy number sequences.

Here, we take a novel approach based on k-mers, short sub-sequences of equal length *k*, generated from whole genome sequencing data of diploid eyebrights (*Euphrasia*), a group of plants which have considerable genome size variation within a ploidy level. We compare k-mer inventories within and between closely related species, and quantify the contribution of different copy number classes to genome size differences. We further assign high-copy number k-mers to specific repeat types as retrieved from the RepeatExplorer2 pipeline.

We find complex patterns of k-mer differences between samples. While all copy number classes contributed to genome size variation, the largest contribution came from repeats with 1000-10,000 genomic copies including the 45S rDNA satellite DNA and, unexpectedly, a repeat associated with an Angela transposable element. We also find size differences in the low-copy number class, likely indicating differences in gene space between our samples.

In this study, we demonstrate that it is possible to pinpoint the sequences causing genome size variation within species without use of a reference genome. Such sequences can serve as targets for future cytogenetic studies. We also show that studies of genome size variation should go beyond repeats and consider the whole genome. To allow future work with other taxonomic groups, we share our analysis pipeline, which is straightforward to run, relying largely on standard GNU command line tools.

## 1 Introduction

Over the past century, cytogeneticists have uncovered various genomic phenomena such as repetitive neocentromers ‘knobs’ (e.g. Creighton and McClintock, 1931), heterochromatin (Heitz, 1928), and B chromosomes (Jones, 1995 and references therein). These are all associated with structural genomic variation, genomic repeats, and they contribute to genome size variation. As recent and ongoing advances in DNA sequencing technology have revolutionised the community’s ability to characterise the genetic variation on the sequence level, it is now possible to study, at unprecedented detail, the sequences underpinning genome size variation within and between closely related species.

Genome size is a trait immediately shaped by structural genomic variation. E.g., a deletion of a part of the genome causes a smaller genome size. Because of the ubiquity in populations of structural genomic variation such as ploidy differences, supernumerary chromosomes, segmental duplications, and other ‘indels’, the assumption of intraspecific genome size variation should be the norm.

However, the magnitude of this variation and whether it can be detected by methods such as microdensitometry or flow cytometry has been subject to debate, and some older reports have been refuted (Greilhuber, 2005; Suda and Leitch, 2010). Nevertheless, studies following best practices and using internal reference standards have revealed genome size variation in numerous species (Achigan-Dako et al., 2008; Šmarda et al., 2010; Díez et al., 2013; Hanušová et al., 2014; Blommaert, 2020).

Between the species of embryophyte plants, genome size shows a staggering 2400-fold variation (Pellicer et al., 2018). Within this range, larger genome size is generally associated with higher proportions of genomic repeats as evidenced by low-pass sequencing studies, although genome repetitiveness deceases somewhat in the species with the largest genomes (Novák et al., 2020a). The repeats accounting for most of the DNA in plant genomes can be classified in two categories: interspersed and tandem (satellite) repeats (Heslop-Harrison and Schwarzacher, 2011) both of which may affect genome evolution in characteristic ways. Interspersed repeats correspond to transposable elements (transposons) which due to their copy-and-paste (or cut-and-paste) nature can insert themselves into distant parts of the genome. Crossing over between such elements can lead to chromosomal rearrangements, associated with DNA loss or duplication, reviewed in Charlesworth et al. (1994). Over evolutionary time, there may be bursts of transposon activity (e.g. Jiménez-Ruiz et al., 2020) possibly triggered by hybridisation (Petit et al., 2010), but short-term change of their copy numbers is usually low. Satellite repeats on the other hand consist of numerous copies arranged in a head-to-tail fashion. Although some satellite repeats are extremely conserved (Abad et al., 1992), they are generally known for rapid changes in copy number and sequence identity between species. This was characterised, in detail, in *Nicotiana* by Kovarik et al. (2008) and Koukalova et al. (2010), and there are numerous other examples for satellite variation between related species (Tek et al., 2005; Ambrozová et al., 2011; Becher et al., 2014; Ávila Robledillo et al., 2020), within populations (Veltsos et al., 2009; Rabanal et al., 2017), and between the sub-genomes of allopolyploids (Heitkam et al., 2020). Satellite copy number has been shown to correlate with genome size for instance in the case of rDNA arrays (Davison et al., 2007; Long et al., 2013) and maize chromosomal knobs (Chia et al., 2012).

Despite the highly advanced state of DNA sequencing and the existence of genome assemblies for many species, it is still challenging to pinpoint the genomic sequences underlying intraspecific genome size variation. This is because structural variation commonly includes genomic repeats, which are often misassembled or missing even in high-quality genome assemblies. Alternative approaches based on low-pass sequencing by design miss low-copy number sequences. In this article, we will demonstrate that comparing the k-mer sets of two individuals allows one to pinpoint in a straightforward way which sequences and genomic copy number classes contribute to genome size differences.

The most familiar representation of an individual-sample k-mer dataset is perhaps a k-mer spectrum as depicted in Fig. 1A. This spectrum plot of the diploid *Euphrasia rostkoviana* shows for each multiplicity level (x axis, number of times a specific k-mer is seen) how many different k-mers there were (y axis). For instance, of k-mers that were observed approximately 130 times there were approximately 3 million different ones (at the tip of the ‘diploid peak’). These k-mers correspond to sequences that were identical between the two genome copies in this diploid individual. There is also a monoploid peak containing sequences present only in one genome only such as caused by heterozygous sites. Repeats are not covered by this plot, which is cropped at multiplicity 200, just above the diploid level. To represent all a genome’s k-mers, an ‘un-cropped’ k-mer spectrum may be plotted with logarithmic axes as in Fig. 1B. Here, the x-axis is labelled with both multiplicity values (black) and the corresponding genomic copy number (grey). The ratio between multiplicity and genomic copy number depends on each individual sample’s sequencing depth. If samples are to be compared, each sample’s multiplicity values must be re-scaled to a common scale, a natural scale being the genomic copy number. To reduce the range of copy number values that are compared, the data may be binned as shown in Fig. 1C, which reduces the number of comparison points to approximately 130 bins (from several 100,000 in Fig. 1B). Because binning is carried out after scaling, a bin number corresponds to the same genomic copy number (range) in all samples.

**Figure 1.**
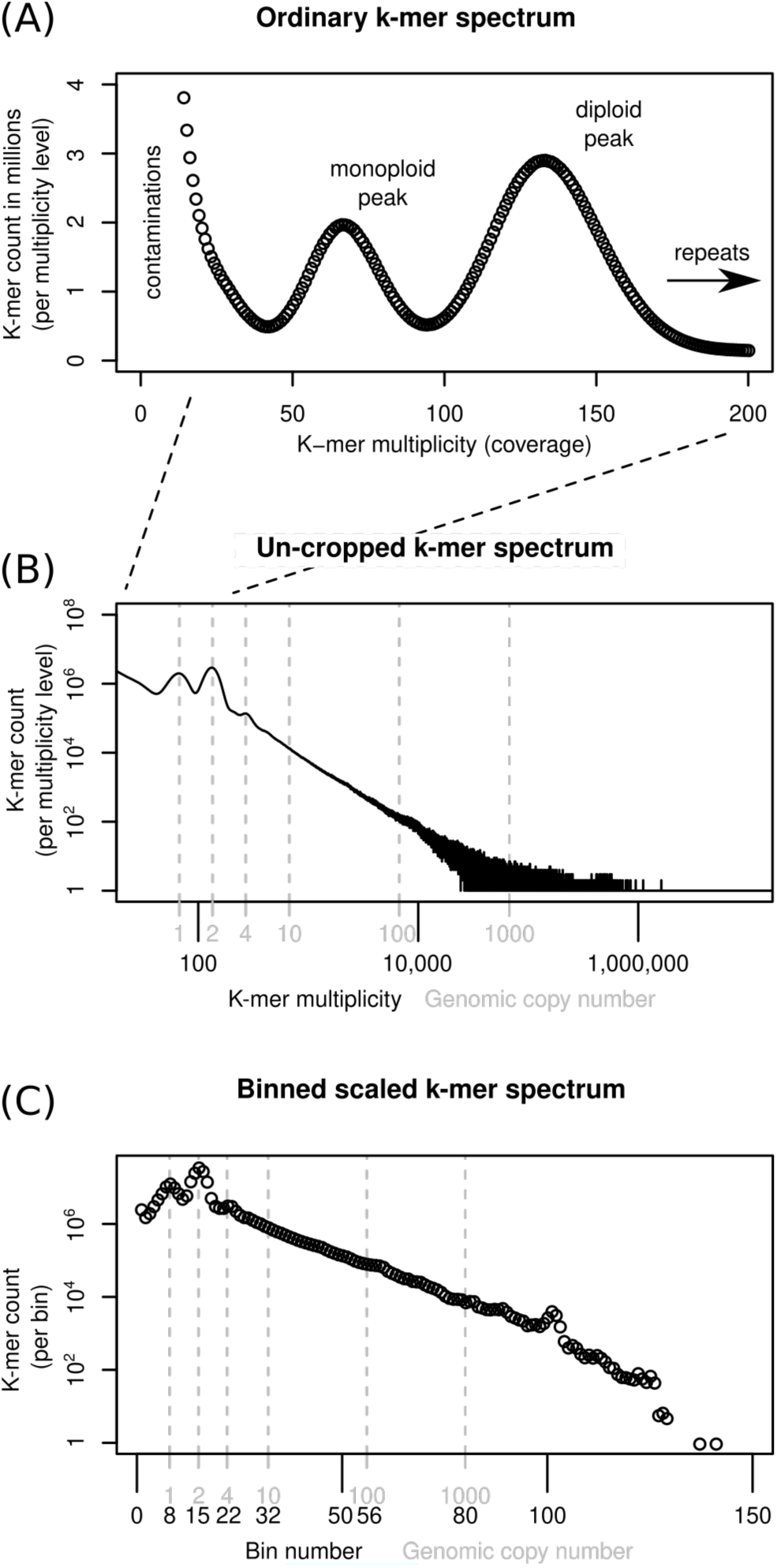
Ways of depicting individual-sample k-mer data sets. Panel (A) shows a k-mer spectrum with linear axes and the multiplicity (x-axis) cropped at 200, excluding k-mers present in genomic repeats. To represent all sample k-mers, the axes may be scaled logarithmically as in (B). To compare samples, the multiplicity values can be scaled and binned (C). See main text for more detail.

Several hypotheses exist as to the sequences causal for genome size differences in closely related species and populations. Here, we investigate three hypotheses, which are not mutually exclusive. (1) Genome size difference may be due to satellite repeats. Satellite repeats are known for their propensity for rapid copy number change as mentioned above and are thus natural ‘suspects’ for causing genome size differences. (2) Differences may be caused by sequences ‘across the board’ – all kinds of sequence proportional to their genomic copy number. Recombination between distant repeat element my cause the duplication, loss, or translocation of larger chromosome fragments resulting in copy number changes of numerous sequences ‘across the board’ (Vitales et al., 2020). (3) Size differences may be due to low-copy number sequences. Numerous pangenome studies have found variation in low-copy number sequences between individuals of the same or closely relates species.

In this study, we use high-coverage shotgun data to investigate the sequences underlying genome size variation in diploid British eyebrights (*Euphrasia* L.) in which we had previously uncovered considerable intraspecific genome size variation (Becher et al., 2021). These diploids from a complex of hybridising taxa, which are not distinguishable by DNA barcoding (Wang et al., 2018) albeit there is some genetic structure congruent with morphological difference as evidenced by AFLPs (French et al., 2008). We intentionally avoid using assembly-based approaches. Instead, we compare genome size and genome composition by means of k-mers allowing us to cover the whole spectrum of genomic repetitiveness classes.

## 2 Materials and Methods

### 2.1 The study system

Eyebrights (*Euphrasia* L., Orobanchaceae) are a genus of facultative hemiparasitic plants with a largely bipolar distribution (Gussarova et al., 2008). All British members of the genus are summer annuals. There are two levels of ploidy know in British eyebrights (*Euphrasia*) – diploid and tetraploid. The diploids tend to have large showy flowers showing a correlation between flower size and degree of outbreeding (French et al., 2005). They carry an indumentum of long glandular hairs and are largely restricted to England and Wales (Metherell and Rumsey, 2018). Tetraploids tend to have smaller flowers, they can have glandular hairs, too, which are then always short, and they occur throughout Britain. Interploidy hybridisation in British eyebrights has been suggested by Peter Yeo, who argued that the diploids *E. vigursii* and *E. rivularis* originated from inter-ploidy hybridisation (Yeo, 1956). So far, only one triploid individual has been confirmed by cytogenetics (Yeo, 1954). In this study, we focus on morphological diploids in which we have previously found 1.2-fold genome size variation (Becher et al., 2021).

### 2.2 Sampling and sequencing

To complement previously generated data (An1, Vi, Ro, and Ri1, see Table 1), we collected morphological diploids in the field and stored samples individually in silica gel for desiccation (see Table 1 for details). We used the UK grid reference finder (https://gridreferencefinder.com) to convert all compute a distance matrix between al sample locations. In total, our sampling covered a geographic range of 570 km (Vi-Ro). Where we included multiple individuals per species, each individual came from a different population with the closest pair of samples being Ri1 and Ri2 collect 2.5 km apart (Table 2).

**Table 1.**

| ID | Species | Read length | Ploi* | Cov* | NCBI ID | % het* | GS (Mbp)* | GS Diff<sup>‡</sup> | Platform<sup>†</sup> | Lat/Long | 1C (pg)<sup>‡</sup> |
| --- | --- | --- | --- | --- | --- | --- | --- | --- | --- | --- | --- |
| An1 | *Euphrasia anglica* | 2 x 250bp | 2 | 54 | SAMN14582932 | 0.13 | 999.98 | NA | X | 50.514/-4.113 | 0.51 |
| An2 | *Euphrasia anglica* | 2 x 150bp | 2 | 28.5 |  | 0.85 | 989.23 | -10.75 | 6 | 51.845/-4.145 | 0.51 |
| Vi | *Euphrasia vigursii* | 2 x 150bp | 2 | 42.4 | SAMN14582918 | 0.14 | 1055.93 | 55.95 | X | 50.24/-5.381 | 0.54 |
| Ro | *Euphrasia rostkoviana* | 2 x 250bp | 2 | 67.4 | SAMN14582916 | 1.13 | 1227.92 | 227.94 | 6 | 55.058/-2.504 | 0.63 |
| Ri1 | *Euphrasia rivularis* | 2 x 150bp | 2 | 35 | SAMN14582917 | 0.23 | 1126.64 | 126.66 | X | 54.534/-3.192 | 0.58 |
| Ri2 | *Euphrasia rivularis* | 2 x 150bp | 2 | 25.5 |  | 1.41 | 1096.44 | 96.46 | 6 | 54.513/-3.203 | 0.56 |
| Ri3 | *Euphrasia rivularis* | 2 x 150bp | 2 | 20.8 |  | 1.41 | 1104.84 | 104.87 | 6 | 53.082/-4.084 | 0.56 |

\* Ploi - ploidy, Cov - multiplicity of the monoploid k-mer peak, % het - heterozygosity in %, GS - genome size in Mbp, each as inferred using Tetmer
<sup>†</sup> Sequencing platform: X - Illumina HiSeq X, 6 - Illumina NovaSeq 6000
<sup>‡</sup> Difference in Mbp to reference individual An1
<sup>‡</sup> Converted following Doležel et al. (2003)

**Table 2.**
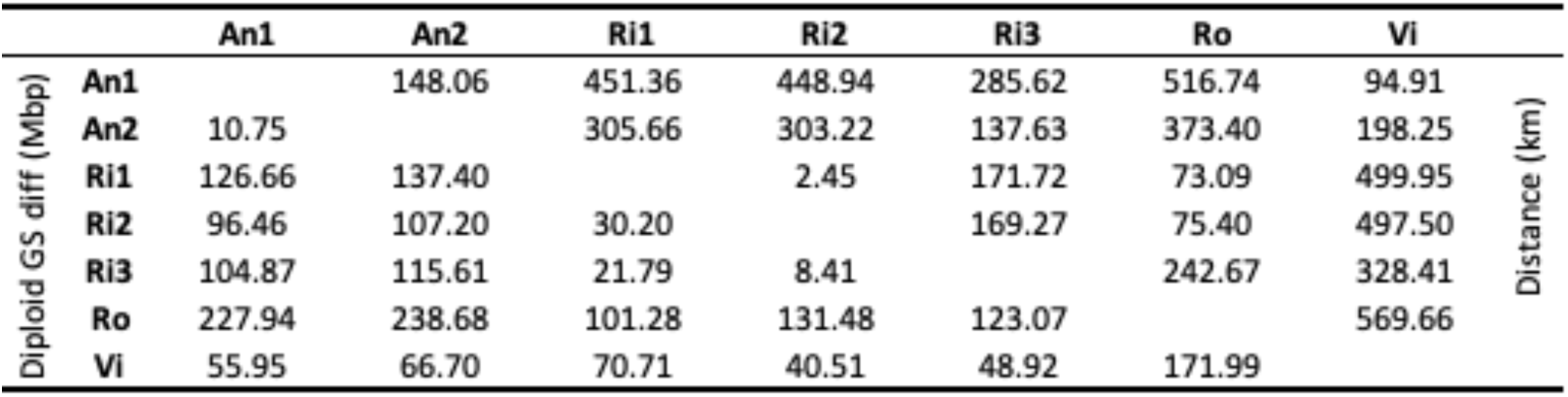
Pairwise diploid genome size differences (below diagonal) and distance between sampling sites (above diagonal) for all sequencing datasets.

We extracted DNA using a DNeasy Plant Mini Kit (Qiagen, Manchester, UK) according to the manufacturer’s instructions. PCR-based libraries were constructed by Edinburgh Genomics, who generated 150-bp paired-end reads on an Illumina NovaSeq6000 instrument.

### 2.3 Handling k-mer data

#### 2.3.1 Generating k-mer data sets and estimating genome sizes

Subsequent to read trimming and filtering with fastp v0.22.0 (Chen et al., 2018) with automatic detection of sequencing adapters in paired-end mode (flag ‘--detect_adapter_for_pe’), we generated k-mer databases for each sample using the software KMC3 (Kokot et al., 2017). Throughout this project, we used 21-mers (k-mers of length 21). In order to remove k-mers of organellar origin, we generated crude *de novo* assemblies of the plastid and mitochondrial genomes using GetOrganelle (Jin et al., 2020) and generated k-mer databases for each organelle. Designed for sequencing data sets, KMC3’s default settings exclude k-mers of multiplicity one, which would likely to be due to sequencing errors. In an assembly, many k-mers will be observed only once. To make all were included, we ran KMC3 with parameter ‘-ci1’. We then used KMC3 to exclude organellar k-mers from each sample database.

For each sample, ee generated three uncropped k-mer spectra (i.e., with the upper multiplicity limit set to 150,000,000, far higher than observed in our data): one for the full (but trimmed and filtered) read data, one with plastid k-mers removed, and one both with plastid and mitochondrial k-mers removed. We profiled these datasets using GenomeScope2, Smudgeplot, and Tetmer.

From these un-cropped, cleaned k-mer spectra we estimated the diploid genome size for each individual as follows. We discarded the portion of each spectrum with multiplicity less than half the individual’s monoploid peak multiplicity, which largely correspond to contamination. For the remaining data, we multiplied the multiplicity and count values. We then took the sum of these products, and divided by the monoploid multiplicity. For conversion to pg, we followed Doležel et al. (2003).

#### 2.3.2 Scaling and binning

To compare between samples the number of k-mers within each frequency (multiplicity) class, we had to scale the multiplicity values of our datasets. We determined for each sample the monoploid (‘haploid’) k-mer multiplicity using the Tetmer app (https://github.com/hannesbecher/shiny-k-mers), and down-scaled the multiplicity values of each k-mer spectrum accordingly so that the resulting spectra had their monoploid peaks at 1 (see Fig. 1B and C). The scaled multiplicity values corresponded to the genome-wide copy number of each k-mer (plus some statistical sampling error caused by shotgun sequencing). However, because each sample had a different monoploid multiplicity, the resulting fraction-valued scaled multiplicity values differed between samples. To compare samples, we binned these scaled multiplicities. Throughout this article, we use the terms scaled (binned) multiplicity and (genomic) copy number interchangeably.

To easily analyse the full range of genomic copy numbers, we decided to use unequal bins, increasing in size in an exponential fashion. We discarded all scaled multiplicities equal to or less than 0.5, because these were likely due to contaminants. We then generated bins (copy number classes) with upper limits 10% larger than their lower limits {(0.5, 0.55], (0.55, 0.605], …, (20.57,22.63], …}. The total number of bins used may differ between samples with the highest bin number corresponding to the highest-copy number k-mer in any dataset. We also generated alphabetically sorted k-mer dumps with KAT3. These are two-column text files of k-mers and their respective multiplicity in a dataset. We scaled and binned these dump files.

#### 2.3.3 Comparing k-mer data sets

Using *E. anglica* (An1) as the reference individual and building on data scaled and binned as described above, we generated two types of sample comparisons: k-mer difference graphs and joint k-mer spectra.

##### 2.3.3.1 Difference graphs

To quantify how much the k-mer differences in each copy number bin contribute to the overall genome size difference between two samples, the per-bin differences are multiplied by the expected copy number of k-mers in each bin. The total genome size difference between two samples can then be obtained by summing over all per-bin products (analogous to computing the genome size from a k-mer spectrum). We generated k-mer difference graphs that indicate the contribution of each copy number bin to the overall genome size difference. This kind of comparison is ignorant of sequence identity. Difference graphs can also be plotted in a cumulative way with the graph’s ‘slope’ indicating the contribution to the genome size difference of any one specific bin. Fig. 2 illustrates for three scenarios how these graphs correspond to the underlying data (here focussing on low-copy number regions).

**Figure 2.**
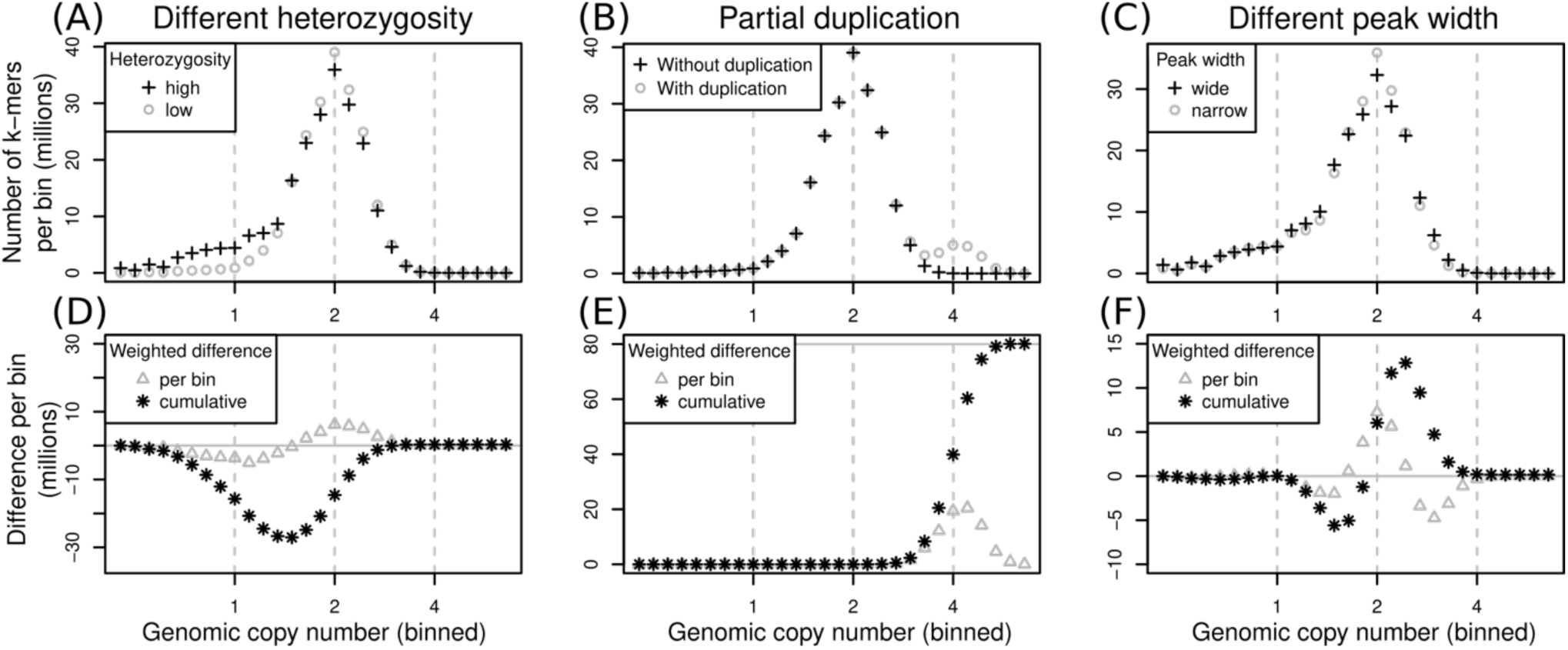
Schematic of pairs of (binned) k-mer spectra (top row) and their corresponding spectrum difference graphs (bottom row). Three different scenarios are shown in columns: (1) two samples of identical genome size with different heterozygosity levels (A and D), (2) two samples where one contains some additional, duplicated sequence (B and E), and (3) two samples with identical sequences but whose k-mer spectra have different peak widths (C and F). Refer to main text for detailed explanations.

The scenarios shown in Fig. 2 are: (1) if one sample has a higher heterozygosity than the other (Fig. 2A) but the samples have identical genome sizes, then the high-heterozygosity sample (crosses) will show a higher 1x peak but a somewhat lower 2x peak than the other sample (circles). The difference graph for this scenario (Fig. 2D) will show two peaks in opposite directions at 1x and 2x (Fig. 2D, triangles). The cumulative difference graph (Fig. 2D, stars) will cross the 1x line with a steep slope indicating a high difference in copy number for 1x k-mers. This is compensated by a steep slope in the opposite direction for 2x k-mer causing a net genome size difference of 0 (vertical grey line). (2) if two samples are identical except for some sequence which is absent in one sample but present at copy number 4 in the other, then one k-mer spectrum will have an additional peak at 4x (Fig. 2B, circles). The corresponding difference graph will show a peak at 4x (Fig. 2E, triangles) and the cumulative difference graph will show a steep slope at 4x leading to a non-zero overall difference (Fig. 2E, stars). (3) different k-mer datasets may have different peak widths even when generated from the same biological sample (technical replicates) depending on the method of library preparation and the sequencing platform chosen. Wider peaks tend to be shallower (Fig. 2C, crosses) than narrow ones (Fig. 2C, circles). This effect may not be obvious in a binned k-mer spectrum, but it does affect difference graphs (Fig. 2F). While not causing the inference of an overall genome size difference, the resulting cumulative difference graph shows a downtick followed by a steep increase crossing x=2 followed by another decrease back to 0 (Fig. 2F, stars). This pattern would be inverted if the samples were swapped.

##### 2.3.3.2 Joint k-mer spectra

A joint k-mer spectrum of two samples is a matrix that shows for each k-mer how often it was observed in each of two datasets. In this way, a joint spectrum is aware of sequence identity. We generated binned joint k-mer spectra by matching up pairs of k-mer dumps (analogous to database joins on the k-mer column). We then scaled and binned the counts in these joins, which reduced the number of count levels from millions to approximately 150 bins. Finally, we counted the number of times that each combination of two bin values occurred, resulting in a three-column table (of count, number of bin in reference, and number of bin in other sample), and we converted this table into a matrix, the binned joint k-mer spectrum. These joint spectra can be visualized as heatmap plots making it possible to show copy number differences between two whole genomes in a single plot.

#### 2.3.4 Contribution of different repeat types

To associate any genomic copy number differences identified using k-mers with specific repeat types, we used the RepeatExplorer2 output of a previous study (Becher et al., 2021) in which we had carried out an analysis of low-pass sequencing data of several diploid and tetraploid British eyebrights. We selected the first 50 repeat super clusters and concatenated, per super cluster, all contributing reads. We then used the program UniqueKMERS (Chen et al., 2021) to extract from each concatenated sequence those k-mers that were unique to the corresponding super cluster, and we turned these into 50 k-mer databases with KMC3. We used these databases to extract from each of the seven high-coverage datasets 50 subsets of repeat k-mers. Finally, we generated joint k-mer spectra for each of these subsets and the corresponding data from reference individual *E. anglica* (An1).

## 3 Results

### 3.1 Genome profiling

Our genome profiling revealed k-mer patterns typical for diploid genomes in all our samples (Table 1). The monoploid k-mer coverage of our datasets ranged from 20.8 in *Euphrasia rivularis* (Ri3) to 67.4 in *E. rostkoviana* (Ro). Per-nucleotide heterozygosity as estimated by Tetmer ranged from 0.13% in *E. anglica* (An1) to 1.41% in *E. rivularis* (Ri2 and Ri3). Samples with very low heterozygosity (such as An1, Vi, and Ri1), containing very few heterozygous k-mer pairs did not have a noticeable ‘AB’ smudge (Supplemental File S1). Smudgeplot incorrectly suggested tetraploidy for these samples, while proposing diploidy for all samples with higher levels of heterozygosity. The spectra’s peak widths (bias parameters) varied considerably between individuals from 0.9 in Ri2 to 2.4 in Vi.

By comparing uncropped k-mer spectra before and after removal of organelle sequences, we could highlight the distributions of organellar k-mers. These had one peak for mitochondrial k-mers (green, Supplemental File S1), but two for plastid k-mers (red, Supplemental File S1). The high multiplicity of these peaks indicating the high copy number of organellar genomes compared to the nuclear ones. The second peak in the plastid-derived k-mers presumably corresponded to the plastid inverted repeat regions. Using un-cropped spectra with organellar k-mers removed, we estimated the genome sizes of our samples to range more than 1.2-fold from 989 Mbp in *E. anglica* (An2) to 1227 Mbp in *E. rostkoviana* (Ro). For comparison, without organellar DNA removed, these estimates were 3.8 to 7.2% higher. Despite our modest sample of seven individuals, the individual genome size estimates showed a clear partitioning by species with ‘species’ accounting for 98.6% of the variation (ANOVA, *F*_3,3_=72.43, *P*=0.0027). Repeating the ANOVA on permuted versions of the dataset showed that this *P*-value and proportion of variance explained are unlikely to occur by chance given a significance cut-off of 5%.

### 3.2 Difference graphs

We generated cumulative k-mer difference graphs for all samples compared to reference individual An1 (Fig. 3). These indicated very similar magnitudes of genome size differences to those obtained from un-binned, un-cropped spectra (Table 2). This suggests that binning, despite reducing the information content of our data, did not bias our inferences.

**Figure 3.**
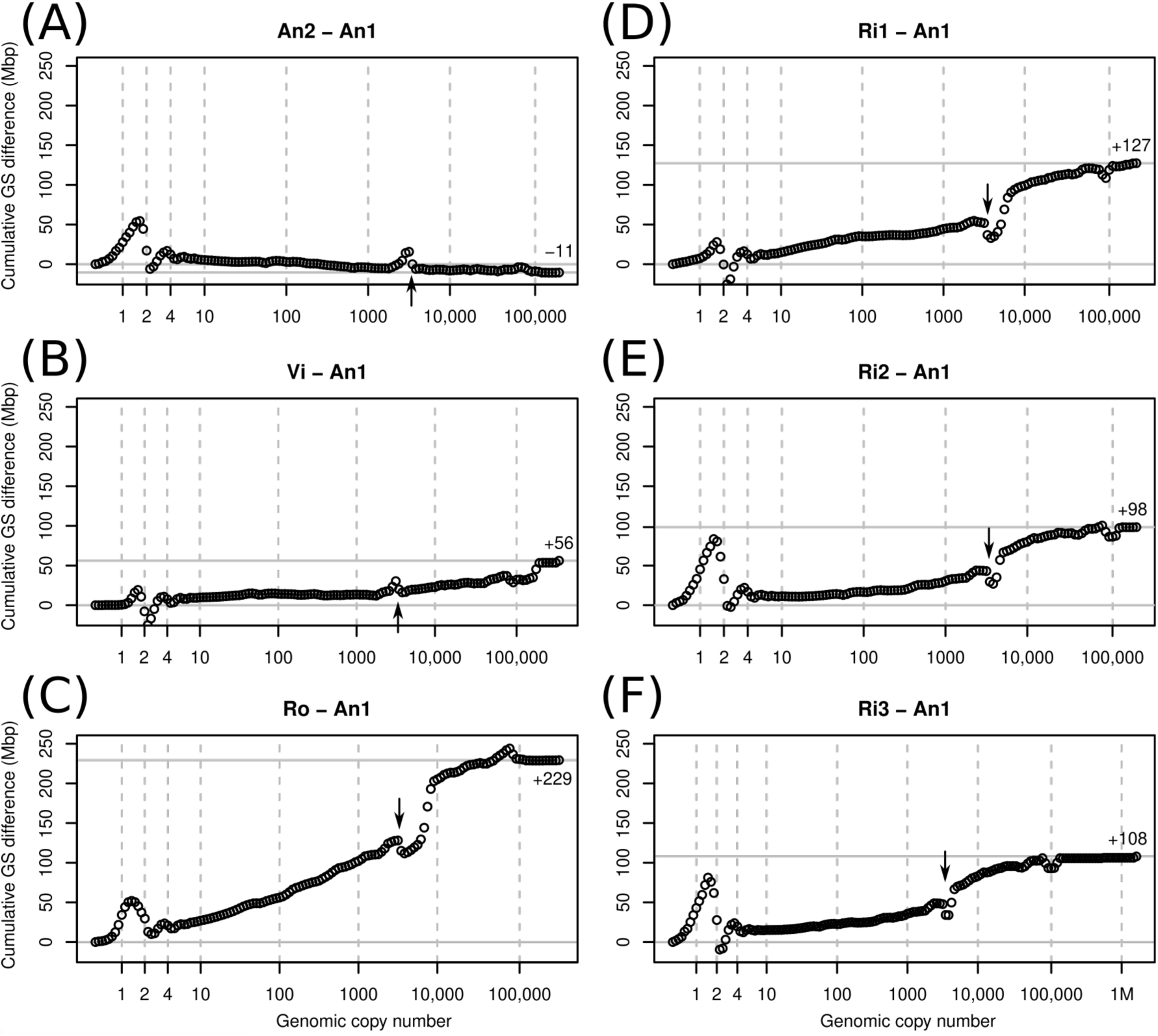
Cumulative k-mer difference graphs of the contributions to genome size differences of genome fractions ordered by increasing repetitiveness for six samples of diploid *Euphrasia* compared to diploid *Euphrasia anglica* (An1). The numbers on the x-axes indicate the genomic copy number bins with 1, 2, and 4 representing haploid, diploid, and ‘duplicated’ sequences. The genome size differences are shown on the y-axes, scaled identically for all graphs. The total genome size difference between the two samples in each graph is indicated at the right-hand side of each plot and by a horizontal grey line. The arrows indicate an anomaly caused by copy number variation of a repeat present in approximately 3000 copies in the reference individual.

The monoploid copy number regions of our cumulative plots are indicated by a vertical dashed line at x=1. These areas of the plots show characteristic differences between low and high-heterozygosity samples. When comparing low-heterozygosity *E. vigursii* (Vi, Fig. 3B) and *E. rivularis* (Ri1, Fig. 3D) to the low-heterozygosity reference individual of *E. anglica* (An1), there were no large differences in heterozygous k-mer counts (which, by definition, have monoploid copy number in diploids) and the curves were flat at x=1. All other samples had higher levels of heterozygosity than the reference individual causing a positive difference in k-mer count leading to a positive slope where the data line intersects with the vertical line at x=1 (Fig. 3A, C, E and F). Again, these are cumulative plots. If the same data were to be plotted per bin as in Fig. 2, positive slopes would be peaks. All samples showed negative slopes where the data line crossed the diploid (x=2) and duplication (x=4) copy number bins. By time the cumulated data series reached x=10 there were no strong up or downticks and all samples had a somewhat higher number of k-mers than the reference individual.

Across the rest of the copy number range, all plots changed largely gradually and nearly monotonically. I.e., across bins, k-mer count differences tended to have the same sign for any individual. An obvious exception from this was a more or less prominent dent in all plots near x=3000 (see arrows in Fig. 3). This pattern is consistent with a repeat of about 3000 copies in the reference sample (An1) and with different copy numbers in the other samples. If a sample contained a lower copy number of this repeat than the reference, then it showed an excess of repeat k-mers at lower copy number followed by a drop at x=3000 as seen in An2 (Fig. 3A) and Vi (Fig. 3B). If, however, a sample contained more copies of this repeat than the reference, then the plots showed a deficiency at x=3000 and a subsequent excess as see in all other samples (Fig. 3C-F). There was a similar, but less pronounced anomaly at approximately x=100,000 in most plots.

### 3.3 Joint k-mer spectra and repeat types

To assess the contribution to genome size differences of individual genomic repeats, we matched up k-mers from our samples with k-mers *Euphrasia*-specific to genomic repeats. We used the 50 largest repeat super clusters identified in a previous study. Collectively, these accounted for approximately 50% of the *Eupharsia* genomes, and the smallest of these superclusters corresponded to a genome proportion of approximately 0.06%. Across samples, the variation in k-mers associated with these repeats accounted for 57% to 78% of the genome size differences observed. Because we used only k-mers unique to individual super clusters, this is likely an underestimate. The only exception was the difference between the *E. anglica* individuals (An2-An1) where the difference in repeat-associated k-mers exceeded the overall genome size difference by 9%. The fact that the An2 genome was larger than predicted based on repeat k-mers suggests that it contained an excess of lower-copy number k-mers compared to the reference individual An1.

Plotting joint k-mer spectra as heatmaps (Fig. 4) allowed us to investigate in more detail how k-mer fractions associated with genomic repeats differed between samples. *E. anglica* (An1) served as reference (along the x axis) in all comparisons. Fig. 4A shows the comparison of all genomic k-mers between Ro and An1. The high heterozygosity of sample Ro showed as dark blue colour at y=1 with the highest counts at y=1 and x=2 indicating that most k-mers found at hererozygous sites in Ro are present in two copies in An1. There is no corresponding high density of k-mers at x=1 and y=2, which agrees with our previous finding of An1 being a low-heterozygosity individual. In the higher-copy number (>1000) regions of the plot, high k-mer densities are found above the diagonal line, indicating higher repeat copy numbers in Ro than An1.

**Figure 4.**
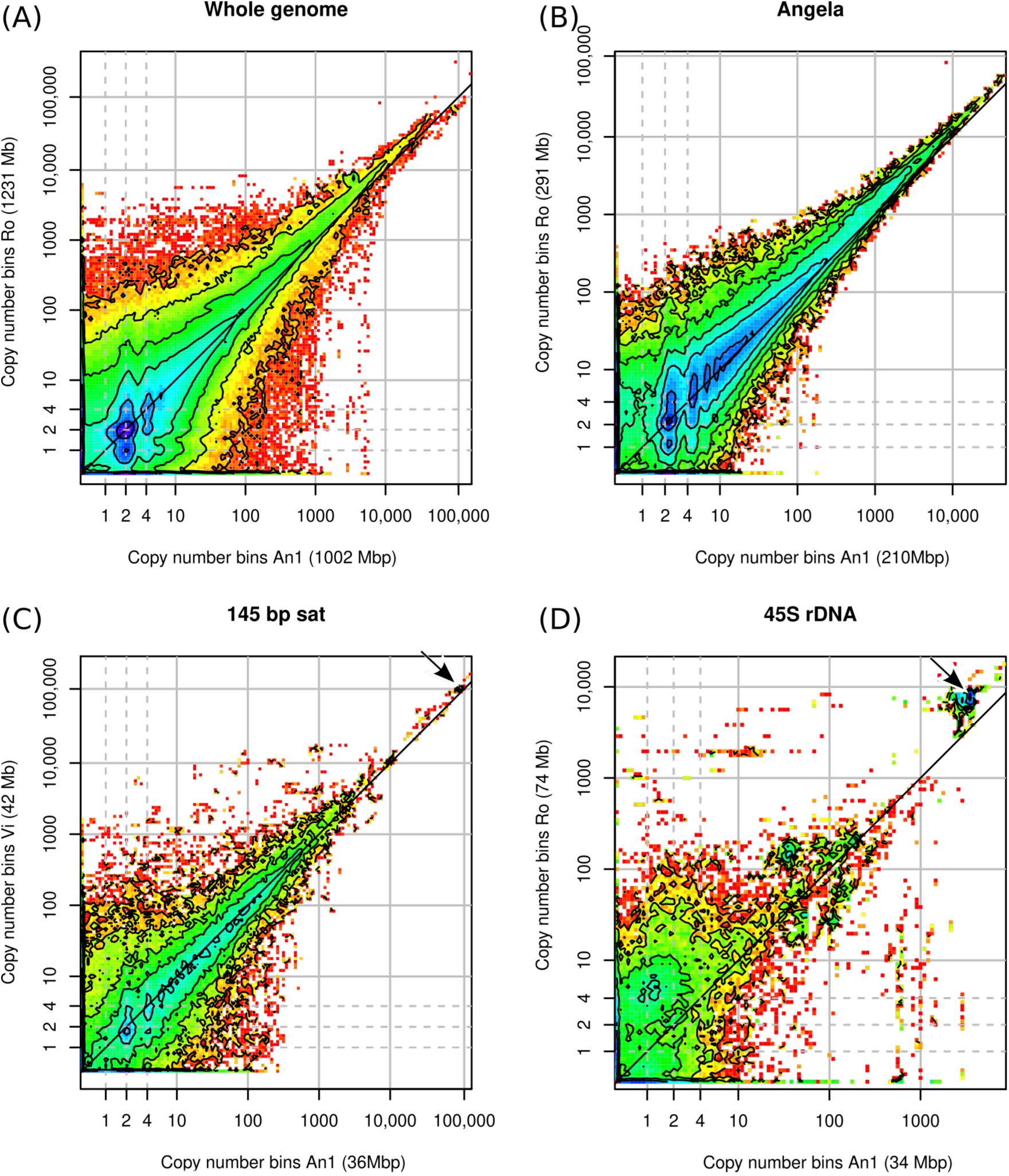
Heatmaps of binned joint k-mer spectra. Copy number bins of the reference individual are shown on the x-axis. The axis labels show in parentheses the contribution of the k-mer fraction depicted to each individual’s overall genome size. The dashed grey lines indicate haploid, diploid and ‘duplicated’ copy numbers. The dark grey diagonal line in each plot indicates the zone where copy numbers are equal between the samples. The arrows in panels (C) and (D) indicate k-mer clusters responsible for the anomalies in Fig. 3.

The repeats with the largest variation between samples in their contribution to genome size were super clusters 1, 4, and 2, which correspond to a Copia transposable element of the family Angela (Fig. 4B), the 45S rDNA, and a 145-bp satellite repeat, respectively. Plotting joint k-mer spectra for individual repeat types, we could match the anomalies seen in the cumulative difference graphs (Fig. 3). The dent at 100,000x corresponds to the 145bp-satellite (Fig. 4C) and the dent at 3000x to the 45S rDNA (Fig. 4D). While the latter two panels contain numerous lower-copy number k-mers in shades of green, yellow, and red, the genome size differences caused by these repeats are accounted for by clusters of high-copy number k-mers located off the diagonal line (indicated by arrows).

### 3.4 The importance of different copy number ranges

To assess which genomic copy number ranges contribute to the overall genome size of an individual, we binned our k-mer spectra even more coarsely. Fig. 5A shows that for all individuals, that the copy number range 0-10 contained the majority of genomic k-mers. The next three copy number ranges, 10-100, 100-1000, and 1000-10,000 contained similar amounts of k-mers, each usually less than half the amount of the 0-10 range. The higher copy number ranges were all smaller. For comparison, we highlighted the contributions to each copy number range of the three largest repeat super clusters 1, 2, and 4 (super cluster 3 corresponded to plastid DNA, which we had removed from our data sets).

**Figure 5.**
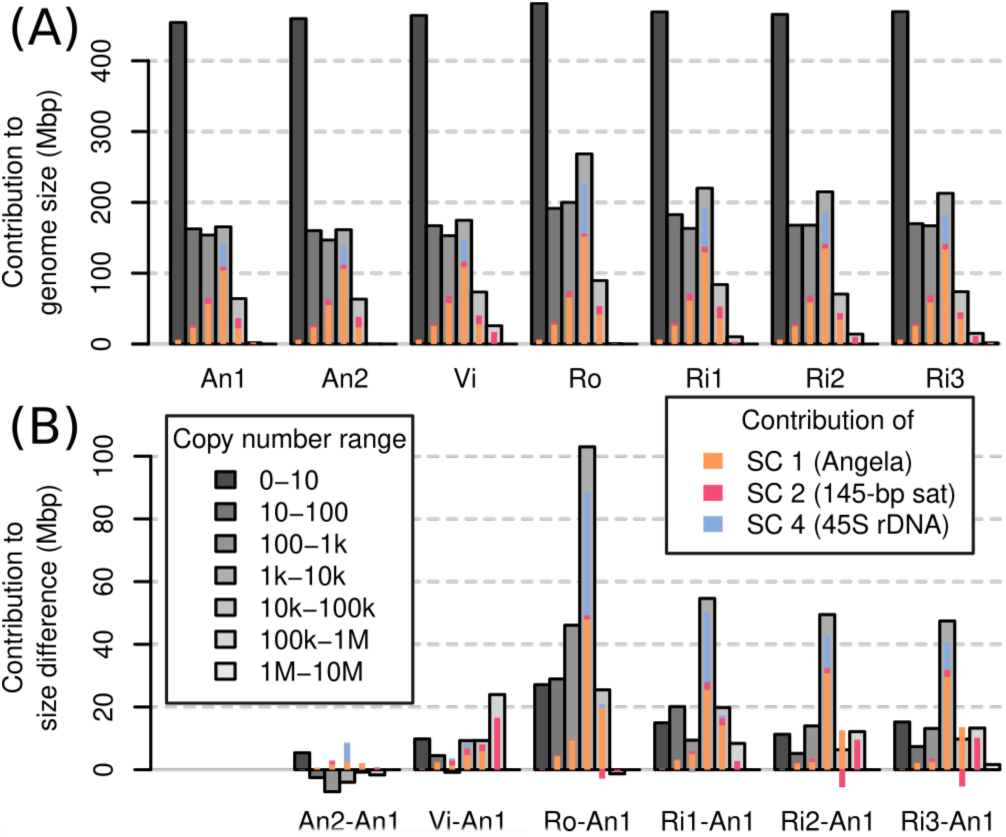
Contribution to overall genome size (A) and genome size differences (B) of genomic copy number ranges. The contributions of repeat super clusters 1, 2, and 4 are indicated in colour.

While the bulk of the samples’ genomes were accounted for by low-copy number sequences (Fig. 5A), we found that the range contributing most to genome size differences was that of 1000-10,000 copies. Most of the differences in this range were driven by sample differences in Angela and 45S rDNA k-mers (Fig. 5B).

## 4 Discussion

In this study, we developed an approach for studying differences in genomic composition within species and between closely related ones, using British eyebrights (*Euphrasia*) as a test case. Rather than using genome assemblies or low-pass sequencing data, we compared the contents of genomes by means of a k-mer approach, which allowed us to inspect the whole range of genomic copy number classes. We found that all copy number classes contributed to genomes size differences with large contributions from a few individual repeats notably including an Angela transposable element. Below, we compare our approach to other existing methods, we critically assess its robustness, and then we turn to what we have learned about eyebright genome evolution.

### 4.1 Comparison to other approaches

The content of two or more genomes may be compared in several ways. Perhaps to most obvious is to use whole-genome alignments, which has been practiced for more than two decades (Chinwalla et al., 2002; Armstrong et al., 2020). Such studies have revealed how genome structure changes over time, for instance following hybridization and whole-genome duplication (Chalhoub et al., 2014). However, most genome assemblies are still not complete, lacking faithful representation of their repetitive sequences. Such sequences are commonly represented in collapsed form or are missing (remaining ‘invisible’) due to the problem of assembling repeats comprising monomers longer than the sequencing read length. Also, genome assemblies usually attempt to represent in one sequence the two (or more) genome copies present in an individual, which may differ in size. Assembly-based approaches are thus unlikely to comprehensively answer the question of genome size differences. Nonetheless, pangenome studies, which compare multiple genomes of closely related species or individuals, have ubiquitously shown that there is structural variation in populations and between closely related species including presence/absence variation of low-copy number sequences (Golicz et al., 2016; Gordon et al., 2017; Hübner et al., 2019).

An alternative approach, focusing only on high-copy number sequences, is the analysis of low-pass genome sequencing data (‘genome skimming’). Because most eukaryote genomes contain more repeats then low-copy number sequences, genome skimming studies can reveal sequences with major contributions to genome size differences. A popular method is RepeatExplorer2 (Novák et al., 2010, 2013, 2020b), which takes a set of short low-pass shotgun sequencing reads, constructs clusters of similar reads, and assembles from these repeat consensus sequences. The repeat clusters are then annotated using a curated database. RepeatExplorer2 can also analyse multi-individual datasets to compare the genome composition of multiple samples, usually of different species. Without the need for a genome assembly, such studies have convincingly shown differences between species in repeat patterns, and plausibly linked these to genome size differences (Ågren et al., 2015; Macas et al., 2015). However, genome skimming studies by design miss single- and low-copy number regions, which also contribute to genome size difference between individuals (Lower et al., 2017).

The approach we chose here may be categorised as a ‘genome profiling’ method, where the properties of genomes are investigated by means of k-mers using moderately high-coverage sequencing data, but in absence of a genome assembly. Other genome profiling methods have been developed to assess assembly completeness (KAT; Mapleson et al., 2016), sequence contamination and heterozygosity (GenomeScope; Vurture et al., 2017), ploidy (Smudgeplot; Ranallo-Benavidez et al., 2020), and to estimate population parameters (Tetmer; Becher et al., 2020). Unlike these single-individual methods, we compared pairs of samples, generating joint k-mer spectra – matrices that simultaneously show the copy number of k-mers in two samples. K-mer multiplicities of individual samples tend to range from one to several millions. Squaring this number, a full joint k-mer spectrum would be too large to handle computationally.

A key aspect of our approach was to bin multiplicity levels, reducing what would be huge un-cropped joint k-mer spectra to matrices of approximately 150×150 bins without losing relevant information. We used these binned joint spectra to compare copy number differences in genome sequences of any copy number, from heterozygous and homozygous single-copy regions (Fig. 4A, blue areas) to satellite repeats (copy number > 100,000, Fig. 4C).

### 4.2 Measuring genome size differences with k-mers

Knowing about the shortcomings of genome assemblies, which tend to be smaller than genomes size estimates obtained by flow cytometry (Bennett et al., 2003), we utilized a k-mer approach in this study. Despite this, we found our bioinformatic genome size estimates were all lower (except for Ro, 1C=0.63 pg) than those we obtained earlier by flow cytometry (Becher et al., 2021), the lowest of which was 1C=0.6 pg. While possible, it seems unlikely that most of our samples truly contained less DNA than all samples analysed in our previous study.

The discrepancy between expected and observed genome size values could not be due to contaminations with non-target DNA, which would have increased, not reduced our estimates. The fact that we removed from our datasets k-mers found in organelle genomes, might wrongly have removed nuclear sequences of organelle origin such as NUMTs or NUPTs, which are known to exist in the family Orobanchaceae (Cusimano and Wicke, 2016), thus biasing downwards our estimates. However these sequences usually account for negligible amounts of the nuclear genome (Hazkani-Covo et al., 2010; Lloyd et al., 2012) + Becher in preparation. Another possibility is that our sequencing data did not contain a faithful representation of the genome contents of our samples. For instance, it is known that Illumina sequencing technologies tend to show a bias against GC-rich sequences.

### 4.3 All frequency classes contribute to eyebright genome size differences

We found that all copy number classes contributed to the genome size differences between our samples. Across most samples, different copy number fractions contributed similar amounts to the overall genome size difference except for the sequences in the copy number fraction 1000-10,000 (Fig. 4B), many of which were 45S rDNA and thus satellite sequences. We also detected a considerable contribution to genome size difference of repeat super cluster 2, which was associated with a 145-bp tandem repeat, possibly a centromeric one, in samples Vi, Ri2, and Ri3 (Fig. 4B). These observations confirm our hypothesis (1) that satellites contribute in a major way to *Euphrasia* genome size differences.

While all copy number classes contributed to the genome size differences, these contributions did not correlate well with the proportion that these copy number class contributed to each genome (compare Fig. 4A and Fig. 4B). For instance, most sequences in all genomes (> 400 Mbp) were low-copy number sequences, which were proportionally underrepresented among the sequences that cause genome size differences. This shows that there was not a per se contribution of all sequences across the board to genome size differences, and we refute our hypothesis (2). However, we cannot exclude the possibility that recombination between distant repeat copies led to copy number changes across numerous sequences. This is because different copy number fractions may not be distributed uniformly along *Euphrasia* chromosomes. For instance, studies on multiple species of grasses have revealed that genomic repeats and single-copy sequence tend to be located in different regions of the chromosomes (Barakat et al., 1998) and it has been shown the gene density in bread wheat increases along chromosomes with increasing distance from centromeres (Akhunov et al., 2003). Although this pattern is not universal (Lang et al., 2018), if it was to hold in *Euphrasia*, structural variation caused by recombination between transposable elements might affect repeat sequences disproportionally more than low-copy number sequences.

Finally, all samples contained more low-copy DNA (copy number < 10) then the reference individual *E. anglica* (An1), ranging from an additional 5 to 27 Mbp at the diploid level. Although this is modest compared to the overall genome size differences between samples, it shows that there is a considerable contribution to genome size differences from low-copy number sequences, which confirms our hypothesis (3). This finding also calls for a *Euphrasia* pangenome study to assess the differences in gene space between *Euphrasia* individuals, which we currently working on.

### 4.4 Genome comparisons and our understanding of diploid British *Euphrasia*

British *Euphrasia* have become known for their taxonomic complexity. While the diploids are largely morphologically distinct from one another (although numerous diploid hybrid combinations are known), they cannot be distinguished reliably by ITS or plastid barcoding (Wang et al., 2018), raising the question whether they are genetically distinct. Adding to this doubt, we have also recently uncovered considerable intra and interspecific genome size variation within *Euphrasia* ploidy levels and showed that ‘population’ is a far better predictor of an individual’s genome size than ‘species’ (Becher et al., 2021). As such, our current working hypothesis has been that *Euphrasia* species may not show genome-wide differentiation, and instead species differences may be maintained by few genomic regions under strong selection while the rest of the genome experiences homogenising gene flow.

These previous findings contrast with our results here, which indicated that genome size is predicted well by morphological species identity and that there are considerable copy number differences in Angela transposable elements between species. Transposable elements are generally thought to show lower rates of copy number change than other genomic repeats and they tend to be dispersed throughout genomes. Divergence in TE copy number might thus indicate genome-wide divergence between the diploid species of British *Euphrasia*. This divergence may not show in the ITS sequences, which due to their repetitive nature tend to show a different turnover behaviour than other nuclear loci. A possible genetic divergence between species may also be missed when analysing plastid sequences, which tend to have lower substitution rates and effective population sizes and thus may not show divergence (Ennos et al., 1999). Introgression (or ‘capture’) of plastid genomes is another increasingly reported phenomenon, which might conceal any existing differentiation in the nuclear genomes. Being mindful of our sampling design, this may be seen as further evidence for diploid British Euphrasia being more distinct species than their tetraploid relatives (French et al., 2008).

## 6 Acknowledgments

We thank the members of the University of Edinburgh’s Genetics Journal Club for feedback on the project. We thank Chay Graham, Kamil Jaron, and Lucía Campos-Dominguez for comments on an earlier version of the manuscript. We also thank Edinburgh Genomics for generating Illumina sequencing data. We thank Chris Metherell for sample identification.

