## Supplemental File S1 for "Measuring the invisible – The sequences causal of genome size differences in eyebrights (*Euphrasia*) revealed by k-mers"

### *Supplementary Material*

#### 1 Supplementary Data

##### 1.1 Genome profiling

All k-mer analyses were carried out with *k*=21.

###### 1.1.1 An1 – *Euphrasia anglica*

Tetmer plot

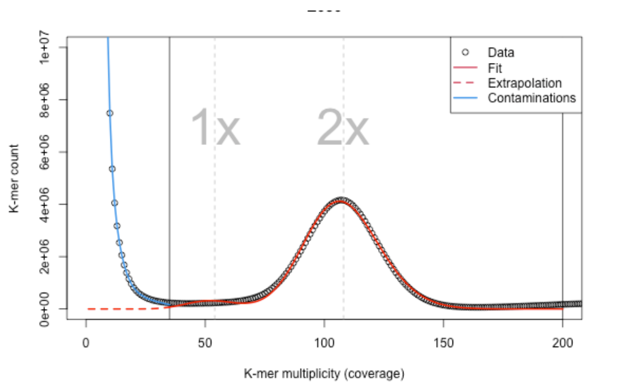


Tetmer fit

```
DIPLOID MODEL, AUTO FITTED
         k-mer length: 21
  monoploid k-mer cov: 54
         θ per k-mer: 0.027
     θ per nucleotide: 0.00128
     non-rep GS (Mbp): 165.5
    bias (peak width): 1.3

STARTING RANGES (MIN MAX)
  monoploid k-mer cov: 10 61
    log10 θ per k-mer: -2 0.6
     non-rep GS (Mbp): 1 2000
    bias (peak width): 0.1 3
              x range: 35 200
```

Smudge plot

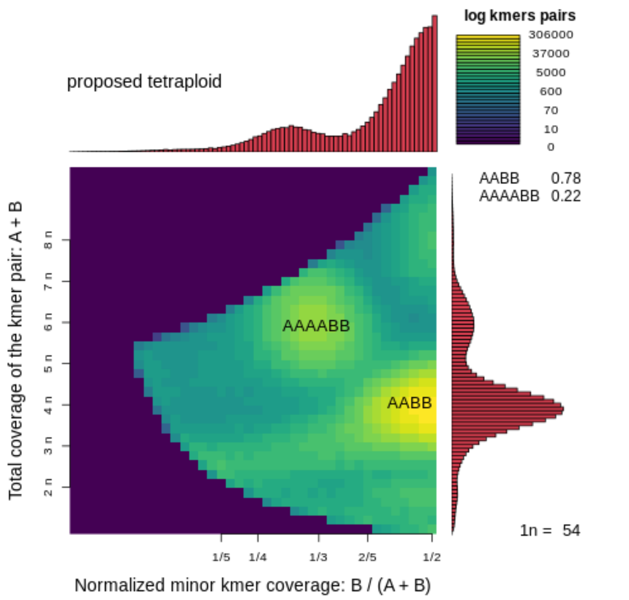


Full k-mer spectrum (axes log-scaled)

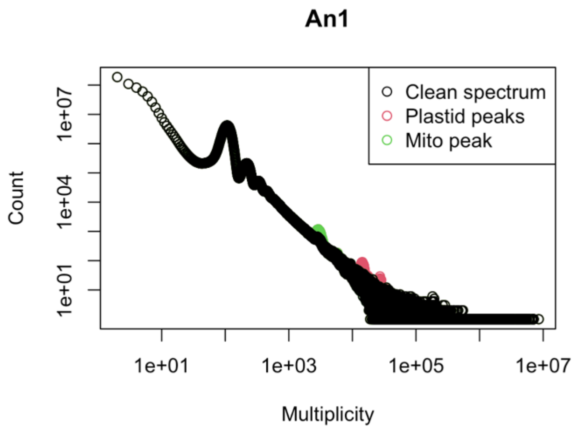

##### 1.1.2 E031 – *Euphrasia vigursii*

**Tetmer plot**

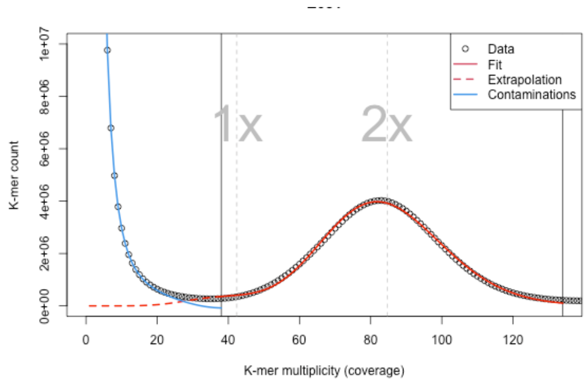


**Tetmer fit**

```
DIPLOID MODEL, AUTO FITTED
         k-mer length: 21
  monoploid k-mer cov: 42.4
          θ per k-mer: 0.0297
     θ per nucleotide: 0.00142
     non-rep GS (Mbp): 171.1
    bias (peak width): 2.4

STARTING RANGES (MIN MAX)
  monoploid k-mer cov: 10 63
    log10 θ per k-mer: -2 0.6
     non-rep GS (Mbp): 1 2000
    bias (peak width): 0.1 3
              x range: 38 134
```

**Smudge plot**

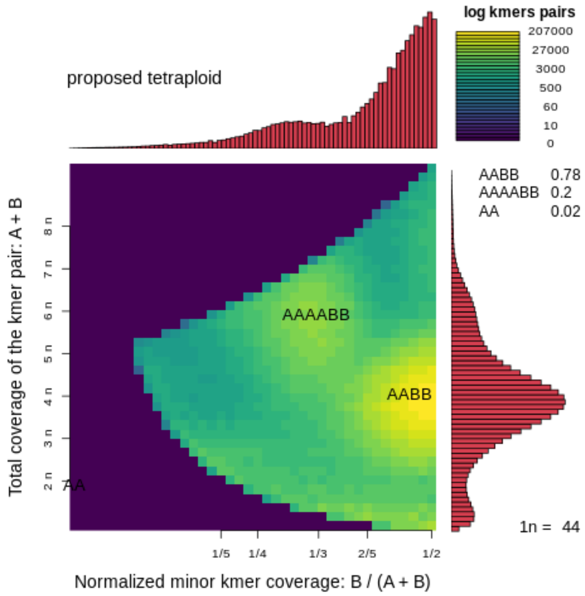


**Full k-mer spectrum (axes log-scaled)**

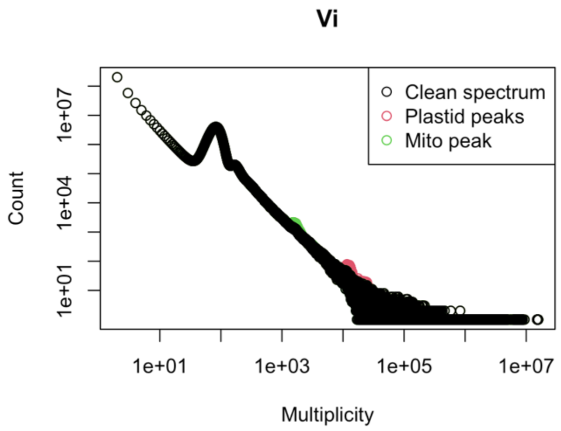

##### 1.1.3 E032 – *Euphrasia rivularis*

**Tetmer plot**

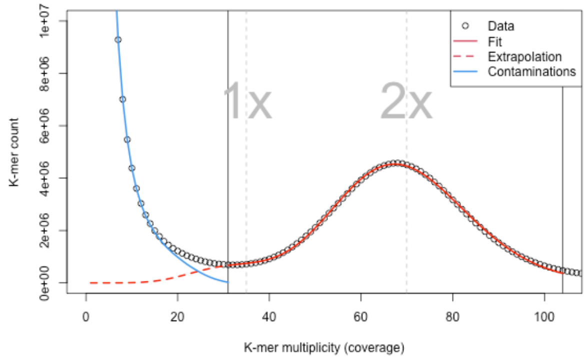


**Tetmer fit**

```
DIPLOID MODEL, AUTO FITTED
         k-mer length: 21
  monoploid k-mer cov: 35
        θ per k-mer: 0.049
     θ per nucleotide: 0.00234
     non-rep GS (Mbp): 170.9
    bias (peak width): 2

STARTING RANGES (MIN MAX)
  monoploid k-mer cov: 10 36
    log10 θ per k-mer: -2 0.6
     non-rep GS (Mbp): 1 2000
    bias (peak width): 0.1 3
              x range: 31 104
```

**Smudge plot**

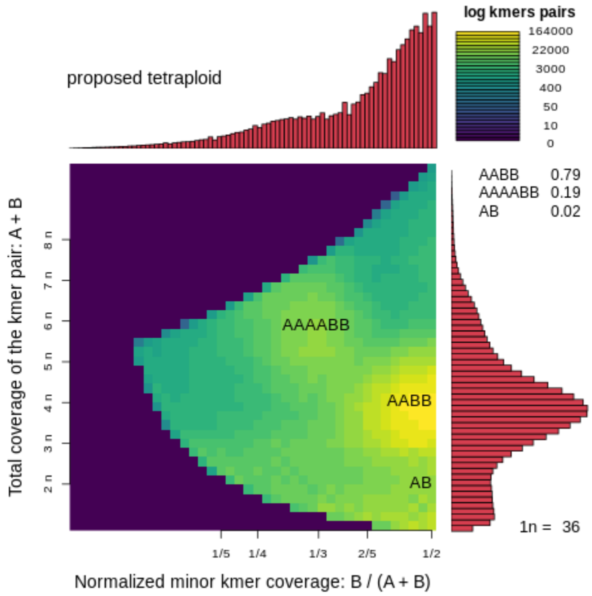


**Full k-mer spectrum (axes log-scaled)**

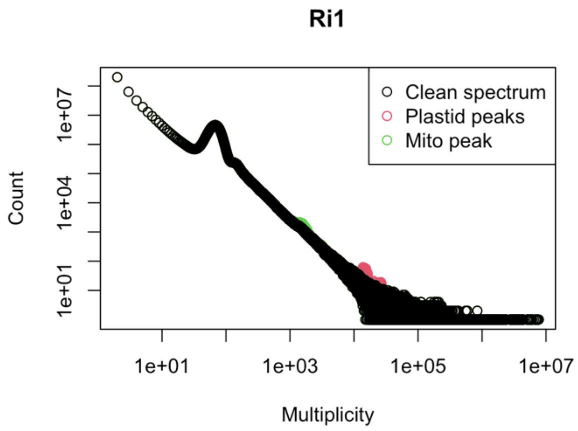

##### 1.1.4 E040 – *Euphrasia rostkoviana*

Tetmer plot

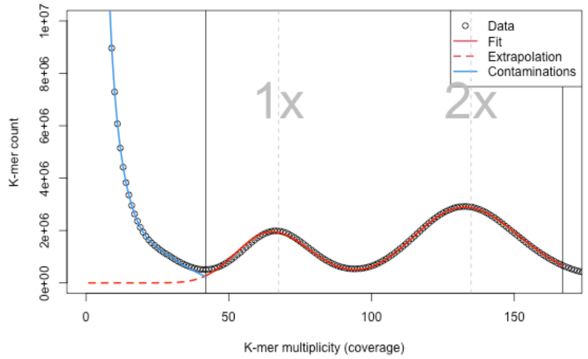


Tetmer fit

```
DIPLOID MODEL, AUTO FITTED
         k-mer length: 21
  monoploid k-mer cov: 67.4
          θ per k-mer: 0.2373
     θ per nucleotide: 0.0113
     non-rep GS (Mbp): 167.8
    bias (peak width): 1.6

STARTING RANGES (MIN MAX)
  monoploid k-mer cov: 10 100
    log10 θ per k-mer: -2 0.6
     non-rep GS (Mbp): 1 2000
    bias (peak width): 0.1 3
              x range: 42 167
```

Smudge plot

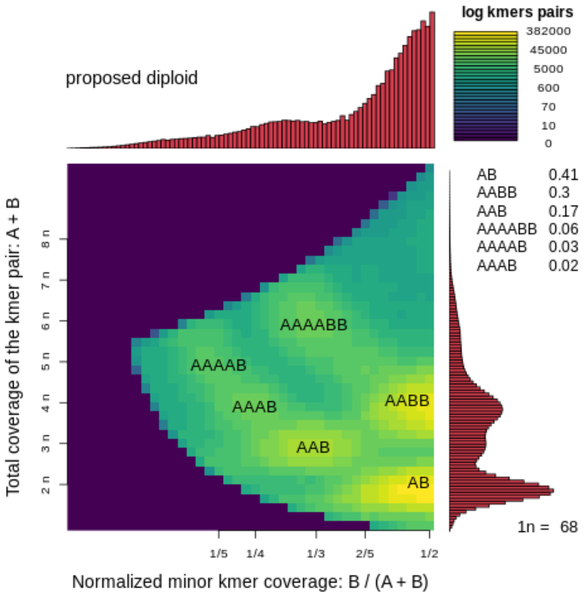


Full k-mer spectrum (axes log-scaled)

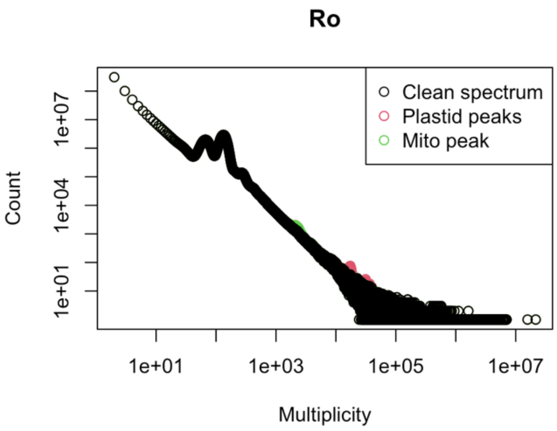

##### 1.1.5 An2 – *Euphrasia anglica*

Tetmer plot

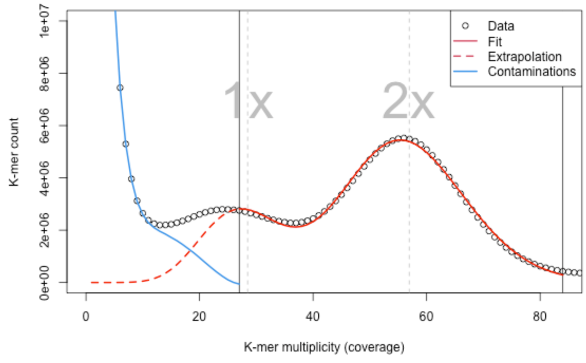


Tetmer fit

```
DIPLOID MODEL, AUTO FITTED
         k-mer length: 21
  monoploid k-mer cov: 28.5
        θ per k-mer: 0.1794
     θ per nucleotide: 0.00854
     non-rep GS (Mbp): 169.2
    bias (peak width): 1

STARTING RANGES (MIN MAX)
  monoploid k-mer cov: 10 44
    log10 θ per k-mer: -2 0.6
     non-rep GS (Mbp): 1 2000
    bias (peak width): 0.1 3
              x range: 27 84
```

Smudge plot

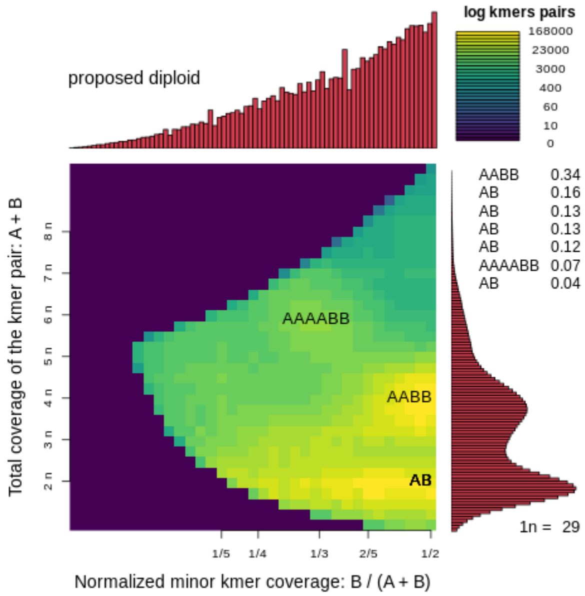


Full k-mer spectrum (axes log-scaled)

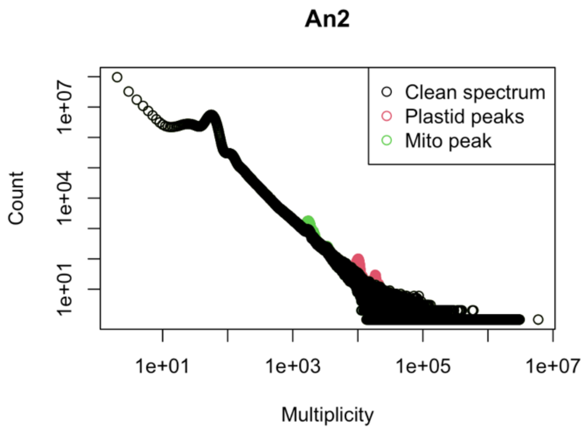

##### 1.1.6 Ri2 – *Euphrasia rivularis*

**Tetmer plot**

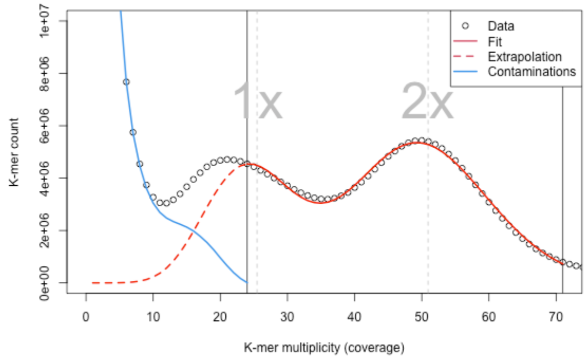


**Tetmer fit**

```
DIPLOID MODEL, AUTO FITTED
         k-mer length: 21
  monoploid k-mer cov: 25.5
          θ per k-mer: 0.2964
      θ per nucleotide: 0.01411
      non-rep GS (Mbp): 169.4
    bias (peak width): 0.9

STARTING RANGES (MIN MAX)
  monoploid k-mer cov: 10 41
    log10 θ per k-mer: -2 0.6
      non-rep GS (Mbp): 1 2000
    bias (peak width): 0.1 3
              x range: 24 71
```

**Smudge plot**

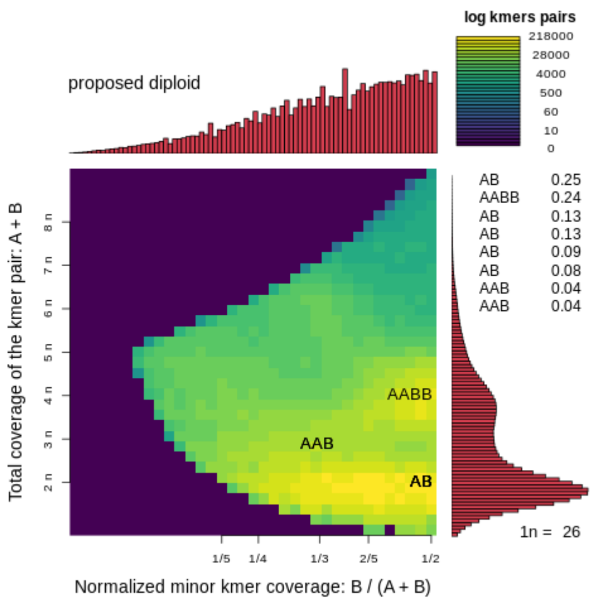


**Full k-mer spectrum (axes log-scaled)**

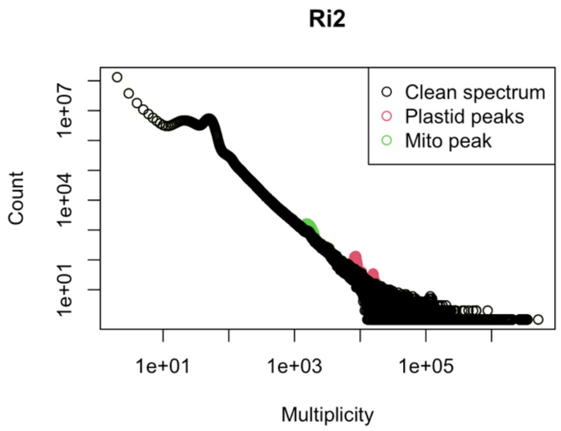

##### 1.1.7 **Ri3 – *Euphrasia rivularis***

Tetmer plot

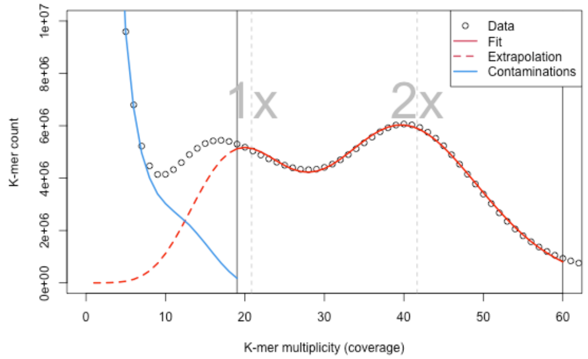


Tetmer fit

```
DIPLOID MODEL, AUTO FITTED
        k-mer length: 21
  monoploid k-mer cov: 20.8
          θ per k-mer: 0.297
      θ per nucleotide: 0.01414
      non-rep GS (Mbp): 171.5
     bias (peak width): 1

STARTING RANGES (MIN MAX)
  monoploid k-mer cov: 10 31
     log10 θ per k-mer: -2 0.6
      non-rep GS (Mbp): 1 2000
     bias (peak width): 0.1 3
              x range: 19 60
```

Smudge plot

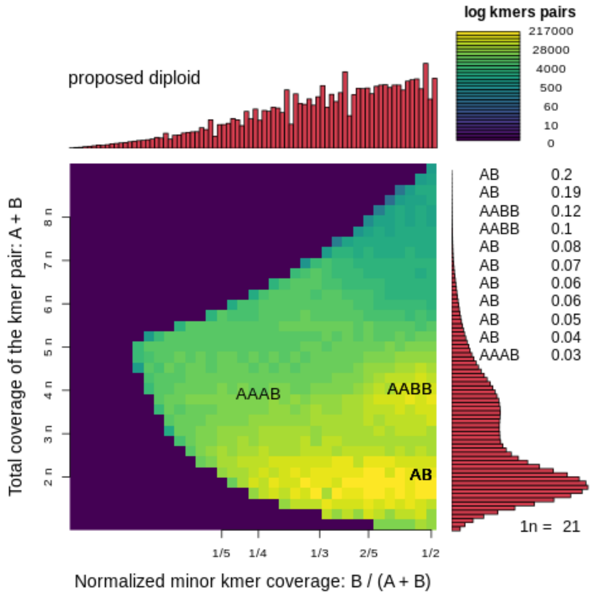


Full k-mer spectrum (axes log-scaled)

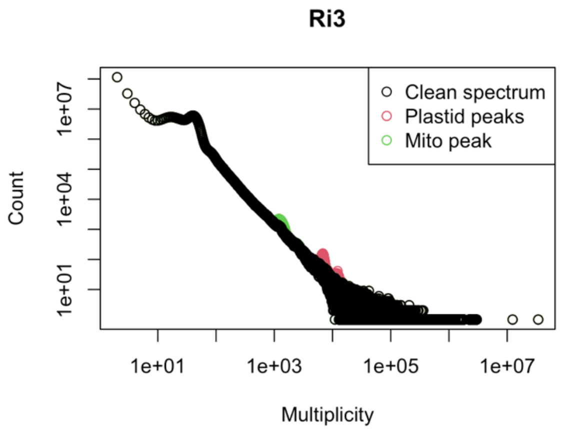
